# A Common Pathogenic Founder Variant in Rwandan Breast Cancer Cases

**DOI:** 10.64898/2026.05.26.727861

**Authors:** Achille VC. Manirakiza, Shakuntala Baichoo, Annette Uwineza, Damas Dukundane, Eulade Rugengamanzi, Janviere Mutamuliza, Allison MO Niragira, Raissa Muvunyi, Jeffrey Besada, Sarah M. Nielsen, Brianna Bucknor, Diane Koeller, Caroline Andrews, Leon Mutesa, Temidayo Fadelu, Timothy R. Rebbeck

## Abstract

Germline data from African populations remain sparse, limiting characterization of population-specific *BRCA1*/2 pathogenic variants. In a study of 175 Rwandan women with breast cancer, 7 unrelated carriers (4% of cases; 22% of pathogenic variant carriers) harbored the same *BRCA1* frameshift variant, c.4065_4068del (p.Asn1355Lysfs*10), which is extremely rare in gnomAD yet recurrent in European, Asian, and Middle Eastern cohorts. Whole-exome sequencing and haplotype analysis of all 7 carriers revealed a shared ancestral block of approximately 581 kb surrounding the variant, and extended haplotype homozygosity and network analyses confirmed a common founder origin. Coalescent-based age estimation placed the founder event approximately 4,000--10,000 years ago. Comparison with 1000 Genomes Project data showed the founder haplotype is absent or exceedingly rare outside African and South Asian populations. These findings strongly suggest the c.4065_4068del variant as a pre-historical *BRCA1* founder variant in Rwanda, with implications for targeted genetic testing, cascade screening, and cancer prevention in the region.

## Background

Breast cancer incidence is rising in sub-Saharan Africa, with an estimated 200,000 cases diagnosed annually. This increasing burden is accompanied by a disproportionate prevalence of aggressive, early-onset cases, and triple-negative breast cancer (TNBC), one of the most aggressive subtypes. Despite this, hereditary variation, particularly germline genomic data from African populations remain sparse^1–3^. *BRCA1* and BRCA2 pathogenic variants (PVs) account for a substantial fraction of hereditary and early-onset breast cancer globally, but most of what is known about variant spectra, founder effects, and population-specific risks comes from European and Asian ancestry cohorts^4–9^. Recent work from East and Central Africa^10–12^, including Rwanda^13, 14^, has begun to close this gap by applying multigene cancer susceptibility panels, revealing *BRCA1/2* PV prevalences that are at least comparable to, and in some series higher than, those observed in European populations, and showing strong enrichment of BRCA1 PV among women with triple-negative breast cancer. These emerging data underscore the need to characterize recurrent and potentially founder PVs in African populations, both to refine risk estimates and to inform context-appropriate genetic testing strategies.

In a recent Rwandan panel-testing study of 175 women with breast cancer, 32 (18%) carried at least one germline PV in an established cancer susceptibility gene, and 7 of these carriers (22%; 7/175 overall, 4%) harbored the same *BRCA1* frameshift PV, c.4065_4068del (p.Asn1355Lysfs*10)^13^. This four-base-pair deletion in exon 10 induces a frameshift at codon 1355 and a premature termination codon 10 amino acids downstream and is predicted to result in nonsense-mediated decay or a severely truncated *BRCA1* protein, consistent with its classification as pathogenic in ClinVar and OncoKB. The variant is extremely rare in general-population datasets (reported in 2–3 of ∼245,000–250,000 chromosomes in gnomAD) but has been described repeatedly in breast and ovarian cancer families from European^4, 15, 16^, Asian^6, 16, 17^, Middle Eastern^7, 18, 19^, and North African populations^9^, and has been implicated as a founder PV in at least one Pakistani series^8^. The unexpectedly high frequency of c.4065_4068del among otherwise unrelated Rwandan cases, all with similar triple-negative phenotypes in prior work^13^, strongly suggests a founder effect in this population. Systematic study of this PV—its frequency, clinical correlates, and underlying haplotype—therefore offers a tractable opportunity to define a potentially important, population-specific *BRCA1* founder variant in Rwanda, with direct implications for genetic risk assessment, cascade testing, and tailored screening and prevention strategies in the region.

## Methods

### Participants

Collection and assessment of 175 breast cancer cases was described previously^13^. Briefly, female breast cancer (fBC) were included if they were 1) Diagnosed on or before the age of 45 years; 2) Diagnosed under age 51 years with unknown or limited family history; 3) Diagnosed under age 51 years with a second breast cancer diagnosed at any age; 4) Diagnosed under age 51 years with 1 or more close blood relatives with breast, ovarian, pancreatic, or PC at any age, 5) Diagnosed under age 61 years with triple-negative breast cancer, the age cutoff of the triple-negative breast cancer was 61 years, and patients with this histology subtype were recruited based on this age limit); 6) Diagnosed at any age with 1 or more close blood relatives with breast cancer diagnosed under age 51 years or ovarian, pancreatic, or high-risk PC at any age; 7) Diagnosed at any age with 3 or more total diagnoses of breast cancer in patient and/or close relatives. A panel of 84 cancer-predisposition genes was used to conduct clinical-grade genetic testing on all patients. Full-gene sequencing, deletion/duplication analysis, and variant interpretation were performed at Invitae (now part of Labcorp) ®.

### Statistical Methods

#### Study Samples and Variant

Whole-exome sequencing (WES) data from 7 unrelated Rwandan individuals, all heterozygous for a pathogenic *BRCA1* frameshift variant (chr17:43,091,462 A>T, proxy for CTTGA>C), were analyzed to characterize the founder haplotype surrounding this variant. The joint-called VCF (GRCh38) was used as the starting point for all downstream analyses.

#### Haplotype Phasing and Tag SNP Selection

Biallelic SNPs within a 2 Mb window centered on the PV were extracted and filtered for minor allele frequency (MAF) ≥ 5% and genotyping rate ≥ 90%, yielding 1,035 variants. Linkage disequilibrium (LD) pruning (window 50, step 5, r² < 0.2) selected 48 tag SNPs. A proxy biallelic SNP for the PV was appended to the tag panel. Statistical phasing was performed using Beagle 5.4 (27Feb25.75f) with the 1000 Genomes Project (1000G) GRCh38 chr17 phased reference panel and a recombination map. Dense phasing of all 1,035 biallelic variants was also performed for higher-resolution analyses.

#### Shared Haplotype Block Detection

For each pair of samples (21 pairs from C (7,2)), the longest shared haplotype block around the PV was identified by extending outward from the PV position and allowing up to 2% allele mismatches to accommodate statistical phasing errors inherent to small sample sizes. For each pair, all four haplotype combinations were tested, and the combination yielding the longest shared block was retained. Physical block lengths (kb) were converted to genetic distances (cM) using interpolation of the GRCh38 genetic map.

#### Variant Annotation

All variants within the *BRCA1* ±1 Mb window were functionally annotated using SnpEff (v5.4a, database GRCh38.99) to predict variant effects (missense, nonsense, frameshift, splice-site, etc.). Clinical significance was overlaid using SnpSift with the NCBI ClinVar VCF database. Annotated variants were summarized into a table with gene, effect, impact tier (HIGH, MODERATE, LOW, MODIFIER), and ClinVar classification.

#### Population Haplotype Comparison

Haplotypes from the 1000 Genomes Project phase 3 (2,548 individuals, 5 superpopulations: AFR, AMR, EAS, EUR, SAS) were extracted at the 14 Rwandan tag SNP positions that overlapped the 1000G chr17 VCF within the shared block region. For each 1000G haplotype, a similarity score (fraction of matching alleles) to the Rwandan founder haplotype was computed.

#### Extended Haplotype Homozygosity (EHH)

EHH decay was computed using the R package rehh on the densely phased Rwandan VCF and on 1000G population subsets (AFR, EUR, EAS). Because the actual PV position was monomorphic in the Beagle-phased VCF (dropped during imputation), a nearby polymorphic proxy SNP (chr17:43,091,983, 521 bp from PV) was used as the focal marker for EHH and bifurcation analyses.

#### Founder Age Estimation

Two complementary approaches were used to estimate the age of the founder mutation. First, a simple decay estimate was constructed using the 7-way shared haplotype block (the longest contiguous region identical across all 7 PV-carrying haplotypes), the block length was converted from base pairs to cM using genetic map interpolation, and the founder age was estimated as g = 1/(2L), where L is the block length in Morgans. Second, a pairwise estimate was computed. For all 21 sample pairs, the optimal shared block length around the PV was computed (with 2% mismatch tolerance). The mean block length across all pairs was used in the same formula, with 95% confidence intervals from 10,000 bootstrap iterations. A generation time of 28 years was assumed for both methods.

#### MDS Analysis

Rwandan and 1000G haplotypes at overlapping tag SNP positions were merged into a single PLINK dataset, and identity-by-state (IBS) distance was computed. Multidimensional scaling (MDS) was performed to project all individuals into a low-dimensional space, with the 7 Rwandan samples overlaid on the 1000G population structure.

## Results

Table 1 describes the clinical and other information for 7 Rwandan breast cancer cases, who were not known to be related to one another, but were found to carry the c.4065_4068del (p. Asn1355Lysfs*10) PV.

**Figure 1** depicts the pedigrees of the 7 putative founder PV carriers. None of these cases was known to be biologically related to any of the others. Detailed clinical information of the patients is provided in Table 2.

**Figure 1:**
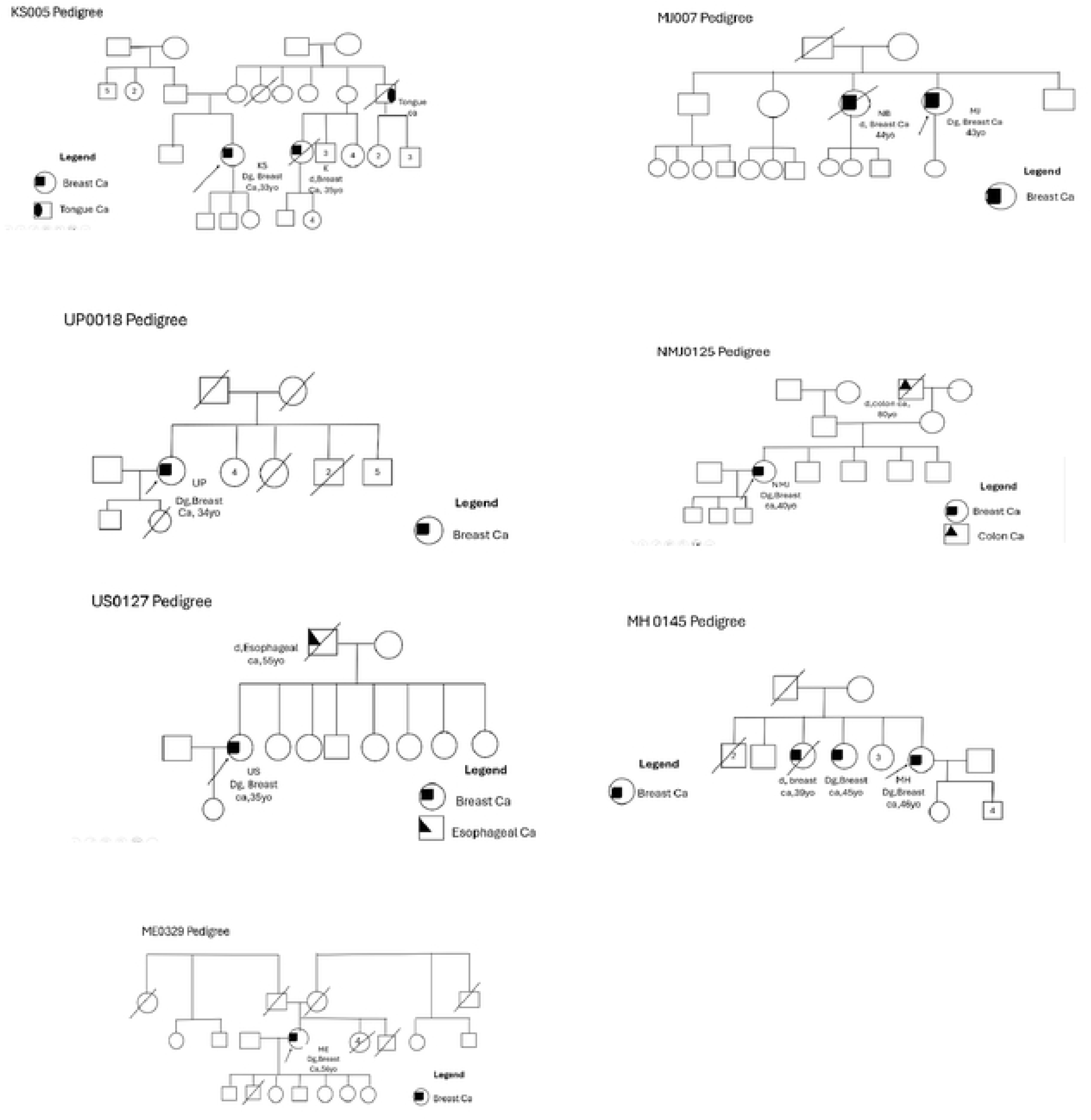
Pedigrees of Putative Founc:ler PV carriers

### Genetic Variant Analysis

A total of 3,865 variants in the *BRCA1* ±1 Mb window was functionally annotated. By SnpEff impact tier: 8 HIGH impact (including the pathogenic frameshift), 48 MODERATE (missense variants), 91 LOW (synonymous), and 3,718 MODIFIER (intronic, intergenic). ClinVar annotation identified 398 variants with clinical classifications: 367 Benign, 16 Likely Benign, 11 Benign/Likely Benign, 2 Variants of Uncertain Significance (VUS), 1 Pathogenic (the known *BRCA1* founder variant), and 1 with conflicting classifications. The predominance of benign variants confirms that the shared haplotype block does not harbor additional pathogenic mutations beyond the known founder variant.

**Figure 2** depicts a large, shared haplotype block that contains the putative founder PV. Pairwise linkage disequilibrium (r²) between the 48 tag SNPs spanning the *BRCA1* ±1 Mb window is displayed as a lower-triangle heatmap. The *BRCA1* gene structure is shown in the top annotation track. The shared haplotype block (green shading, 42.65–43.23 Mb) and the pathogenic variant position (red crosshairs) are marked. A prominent block of high LD (dark red tiles) is visible within the shared region, consistent with the founder haplotype being in strong linkage across this segment.

**Figure 2:**
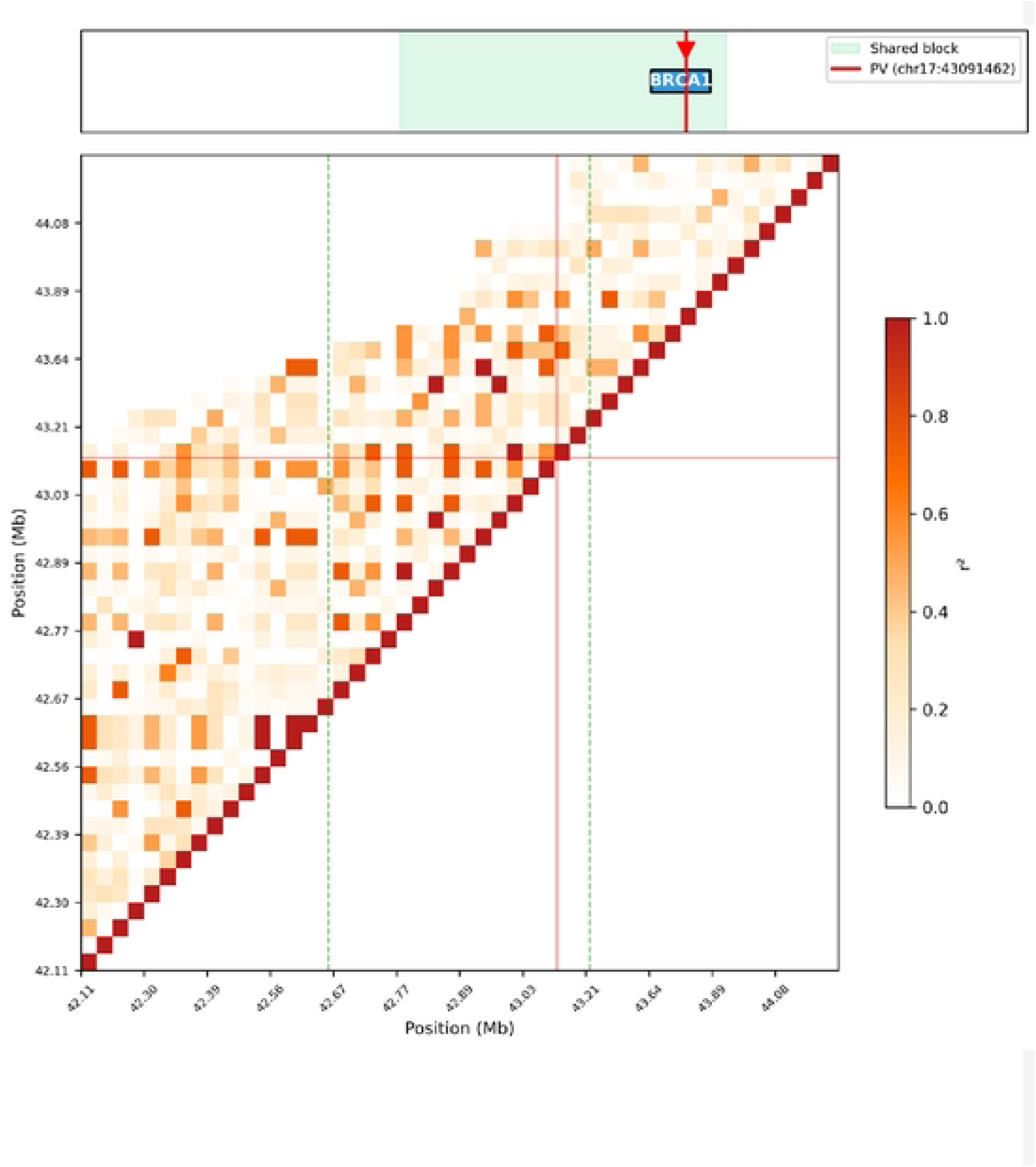
BRCA1. LD structure **With gene** annotation.

**Supplementary Figure 1** depicts the phased haplotype structure of the 7 putative founder PV carriers. ach row represents one phased haplotype (14 total: 7 samples × 2 haplotypes). Blue tiles indicate the reference allele, and red tiles indicate the alternate allele at 1,282 informative variant positions across the ±1 Mb window. The green-shaded region marks the shared haplotype block (∼581 kb). The red dashed line indicates the pathogenic variant at chr17:43,091,462. Within the shared block, a clear pattern of allele identity is visible on the PV-carrying haplotypes, supporting a common founder origin.

**Figure 3** reports haplotype similarity between Rwandan founder and 1000 Genomes populations. Panel A shows violin plots of haplotype similarity scores (fraction of matching alleles at 14 overlapping tag SNPs) between the Rwandan founder haplotype and 5,096 1000G haplotypes, grouped by superpopulation. The red dashed line marks the maximum achievable similarity (13/14 = 0.93). African (AFR) populations show the broadest distribution, with 3 haplotypes reaching the maximum similarity. South Asian (SAS) populations show the highest mean similarity (0.489). Panel B shows the frequency of high-similarity haplotypes at three thresholds (>70%, >80%, >90%). SAS has the highest proportion at >80% (63 haplotypes, 6.4%), while only AFR and SAS populations contain haplotypes at >90% match. The absence of any 100% match in 1000G suggests this specific founder haplotype may be private to or enriched in Rwandan/East African populations.

**Figure 3.**
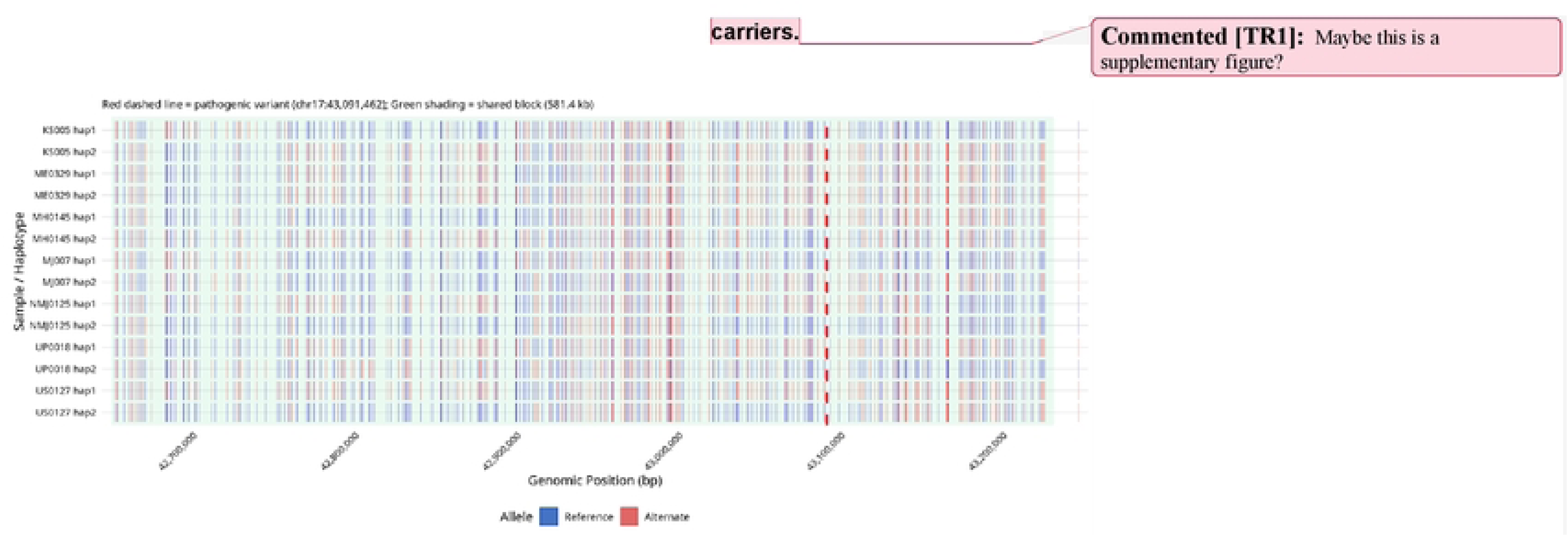
Phased haplotype structure of 7 **Rwandan BRCA1** arriers

**Figure 4a** depicts the extended Haplotype Homozygosity (EHH) decay in Rwandan samples. EHH is plotted for the ancestral (red) and derived (blue) alleles at the nearest polymorphic proxy to the PV in the 7 Rwandan samples (14 haplotypes). The ancestral allele shows a sharp peak of haplotype homozygosity near the PV (reaching 1.0) with rapid decay over ∼10 kb, consistent with a relatively young founder event that has not yet been broken up by recombination in this small sample. The derived allele EHH is near zero across the region, reflecting very low frequency of the derived allele at this specific proxy position. Note: with only n=7, EHH estimates are noisy.

**Figure 4.**
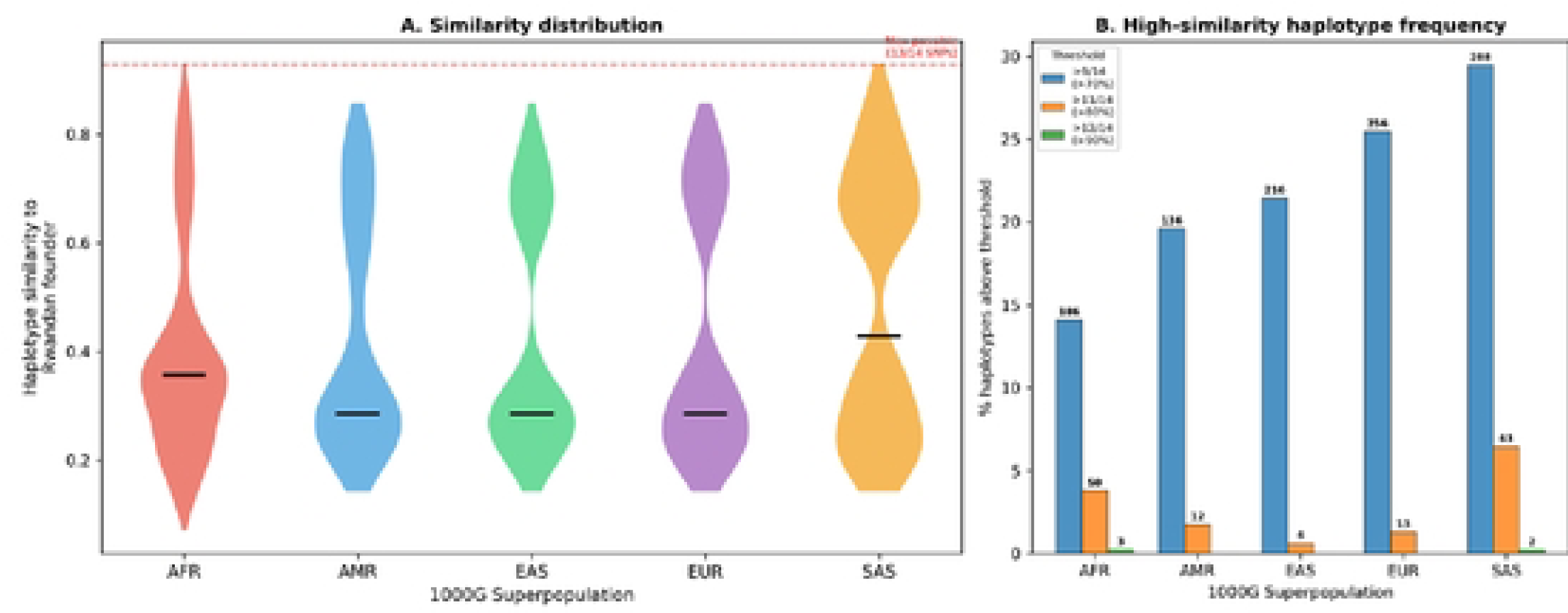
Haplotype similarity between Rwandan founder and 1000 Genomes populations.

EHH decay is shown in **Figure 4b** for three 1000G superpopulations at the nearest SNP to the *BRCA1* PV: AFR (n=660, red), EUR (n=503, purple), and EAS (n=504, green). In all three populations, the derived allele shows a sharp EHH peak centered on the PV position with rapid decay within ∼20 kb. The AFR population shows a broader and more elevated EHH for the ancestral allele, consistent with greater haplotype diversity at this locus in African populations. The sharp EHH peaks in all populations indicate that the variant sits within a region of high haplotype homozygosity, though the pattern does not suggest strong recent positive selection at this locus.

Haplotype bifurcation is shown in **Supplementary Figure 2** for the ancestral (red, 6 haplotypes) and derived (blue, 8 haplotypes) alleles at a polymorphic proxy SNP (chr17:43,091,983, 521 bp from the PV). Each horizontal line represents one haplotype extending outward from the focal SNP in both directions, with branching points indicating where haplotypes diverge. The derived (blue) haplotypes show tight clustering near the focal point with a single long shared tract, consistent with a recent common ancestor. The ancestral (red) haplotypes diverge more rapidly, indicating greater background haplotype diversity. The dashed line marks the actual PV position (chr17:43,091,462). With only 14 haplotypes, the diagram is sparse but clearly shows the extended sharing among PV-associated haplotypes.

**Supplementary Figure 3** shows the pairwise shared haplotype block length distribution. Panel A shows the histogram of pairwise shared block lengths (in cM) across all 21 sample pairs. The distribution is strongly right-skewed: most pairs share short blocks (median = 0.032 cM), while a subset shares much longer blocks (up to 0.33 cM), reflecting heterogeneity in the extent of shared ancestry. Panel B displays block sizes (in kb) for each pair, sorted by length and color-coded (green = above median, red = below). Pairs involving KS005, ME0329, and UP0018 consistently show the longest shared blocks (>400 kb), suggesting these three individuals share the most recent common ancestor for the PV-carrying haplotype. The mean and median lines are annotated directly below the bars.

**Figure 5** depicts the haplotype network of 14 phased *BRCA1* haplotypes. Multidimensional scaling (MDS) of pairwise Hamming distances (computed across 14,489 variants in the shared block region) positions the 14 haplotypes in two dimensions. Minimum spanning tree (MST) edges connect haplotypes, with edge labels indicating the number of SNP differences. Each node is coloured by sample. The three PV-carrying founder haplotypes (KS005 hap2, ME0329 hap2, UP0018 hap1; marked [FOUNDER]) cluster tightly on the right with only 1–2 SNP differences, confirming they carry a virtually identical haplotype inherited from a common ancestor. KS005 hap1 and MJ007 hap1 are completely identical (0 differences), forming a shared non-PV haplotype on the left. The remaining haplotypes show 130–530 SNP differences, representing normal background haplotype diversity.

**Figure 5.**
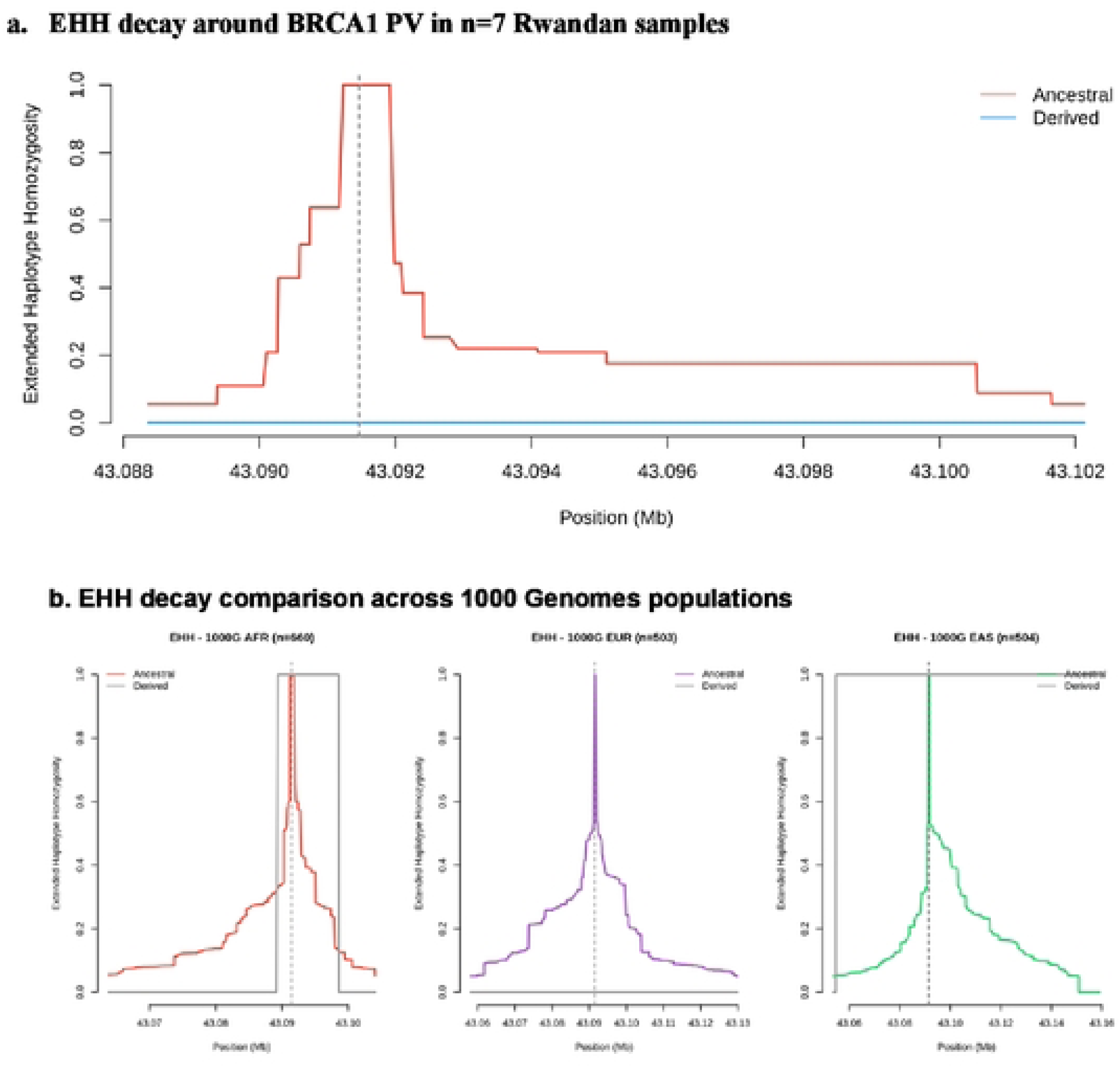
Extended Haplotype Homozygoslty (EHH) decay In Rwandan samples.

**Figure 6.**
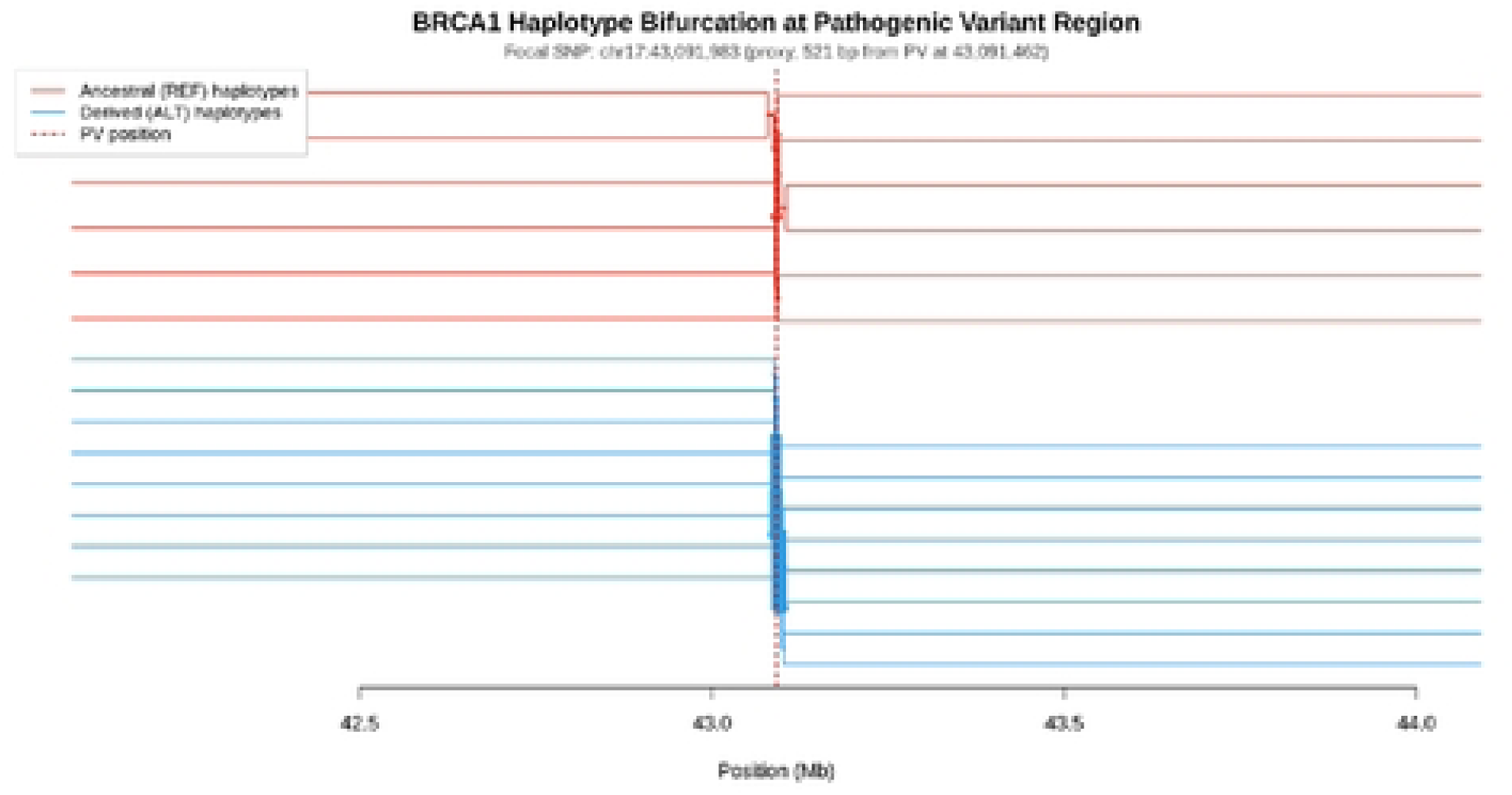
Haplotype bifurcation diagram at the BRCA1 pathogenic variant region.

**Figure 7.**
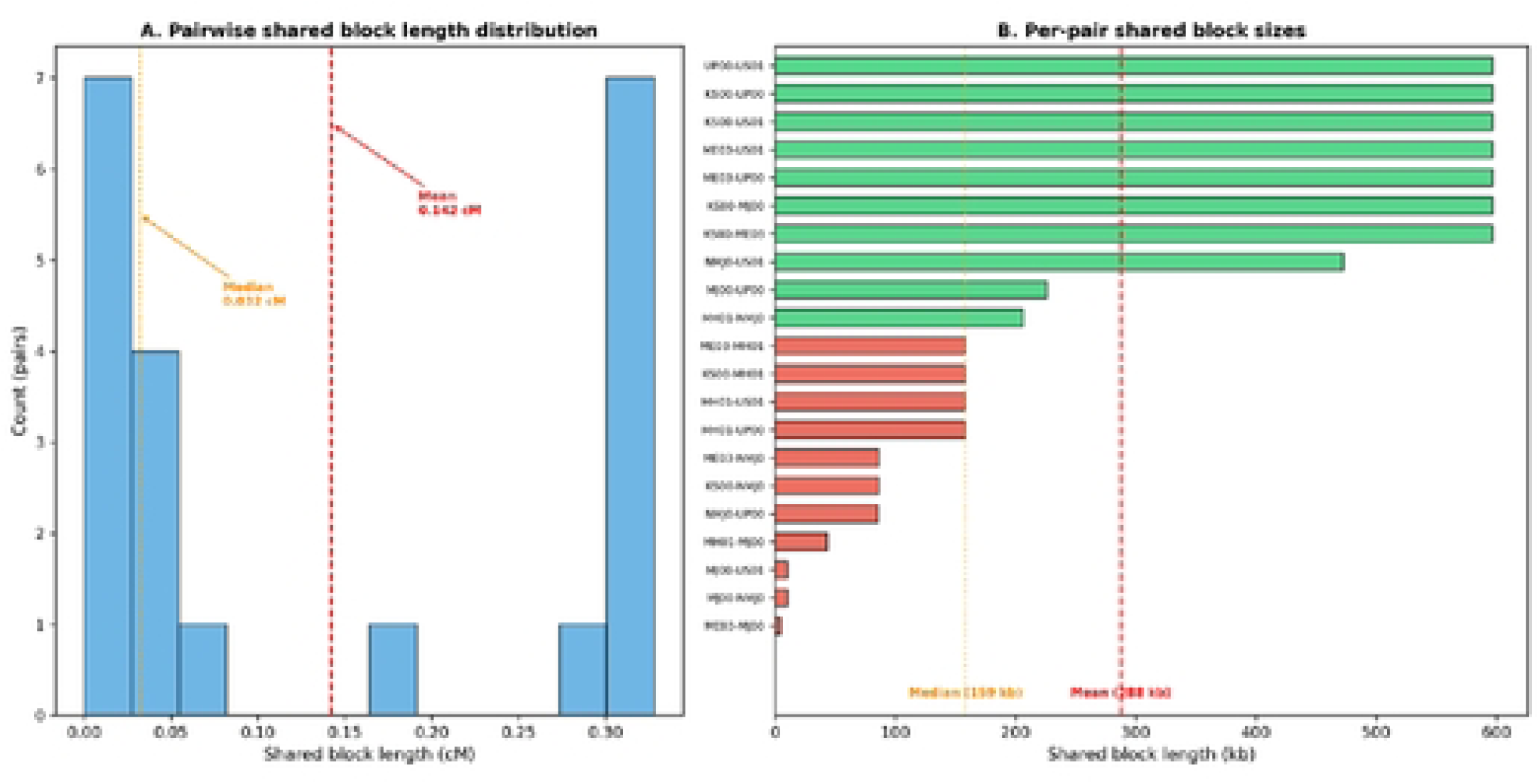
Pairwise shared haplotype block length distribution.

**Figure 8.**
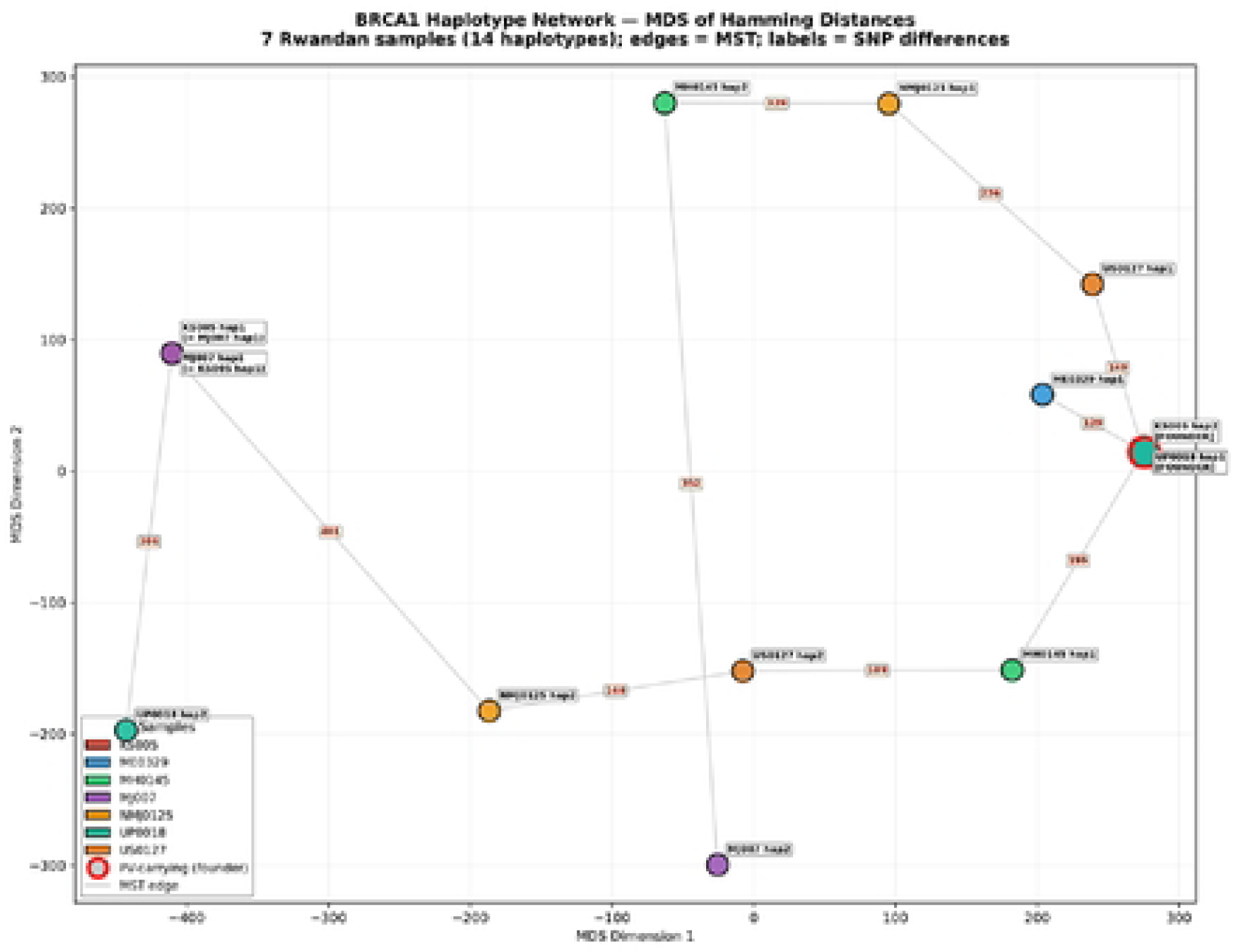
Haplotype network of 14 phased BRCA1 haplotypes.

**Figure 9.**
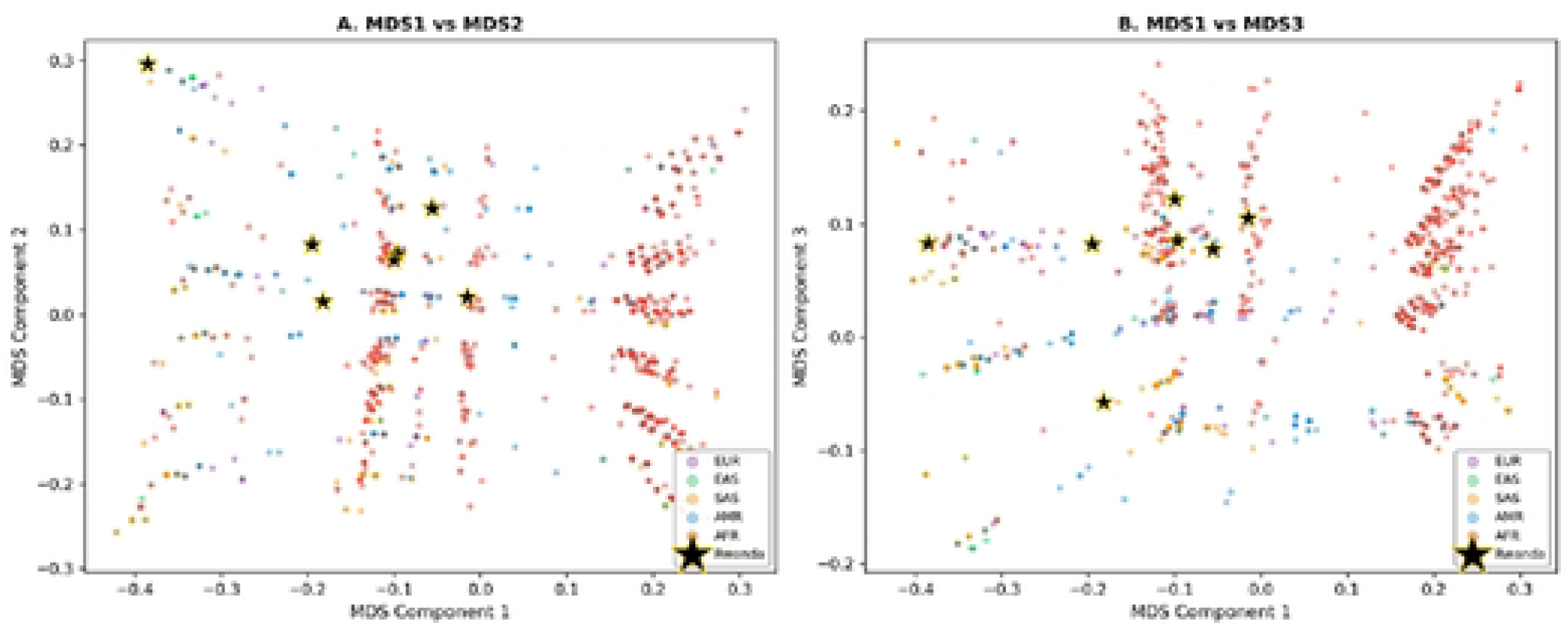
MDS or BRCA1 haplotype sharing: Rwandan vs 1000 Genomes populations.

MDS of *BRCA1* haplotype sharing in Rwandan vs 1000 Genomes populations is shown in **Supplementary Figure 4.** IBS-based MDS projects the 7 Rwandan samples (black stars with gold edges) onto the haplotype structure of 2,548 individuals from the 1000 Genomes Project at 14 overlapping *BRCA1* tag SNP positions. Panel A (MDS1 vs MDS2) and Panel B (MDS1 vs MDS3) show that the Rwandan samples are dispersed primarily within and near the African (AFR, red) cluster, consistent with their population of origin. Some Rwandan individuals overlap with AMR and SAS clusters on certain MDS axes, reflecting the limited resolution of 14 tag SNPs for fine-scale population differentiation. The MDS confirms that the *BRCA1* haplotype structure of Rwandan carriers is broadly African but does not closely match any single 1000G sub-population, consistent with the founder haplotype being locally enriched in the Rwandan population.

### Patient Relatedness

Because the 7 founder PV carriers were not known to be biologically related, we explored relatedness among these individuals. IBS-based MDS projects the 7 Rwandan samples (black stars with gold edges) onto the haplotype structure of 2,548 individuals from the 1000 Genomes Project at 14 overlapping *BRCA1* tag SNP positions. Panel A (MDS1 vs MDS2) and Panel B (MDS1 vs MDS3) show that the Rwandan samples are dispersed primarily within and near the African (AFR, red) cluster, consistent with their population of origin. Some Rwandan individuals overlap with AMR and SAS clusters on certain MDS axes, reflecting the limited resolution of 14 tag SNPs for fine-scale population differentiation. The MDS confirms that the *BRCA1* haplotype structure of Rwandan carriers is broadly African but does not closely match any single 1000G sub-population, consistent with the founder haplotype being locally enriched in the Rwandan population.

### Founder Age Estimation

We used two approaches to estimate the age of the putative founder PV. First, a simple decay-based estimate was applied. The 7-way shared haplotype block spans chr17:42,650,027–43,231,456 (∼581 kb). Using the GRCh38 genetic map, this corresponds to a genetic distance of 68.2375–68.5505 cM, yielding a block length of 0.3129 cM (0.003129 Morgans). Applying the standard decay formula g = 1/(2L), we estimated that the founder arose approximately 160 generations ago, with an estimated age of 4,474 years (assuming 160 generations × 28 yr/gen), or approximately 2,448 BCE, This simple estimate uses the longest contiguous block shared by all 7 carriers and provides a lower bound on the mutation age, since the 7-way block represents the minimum shared region across all haplotypes.

Second, we used a pairwise estimate with bootstrap CI to examine all 21 sample pairs (C(7,2)) using optimal haplotype matching with 2% mismatch tolerance to accommodate phasing errors. The distribution of pairwise shared block lengths was strongly right-skewed. The mean shared block was 0.1424 cM (287.7 kb) (Median shared block: 0.0319 cM; 158.6 kb). Using this approach, we estimated that the PV arose about 351 (95% CI: 242–620) generations ago, with an estimated mean age of ∼9,831 years (95% CI: 6,764–17,357 years). The pairwise estimate yields an older age than the simple decay method because it incorporates shorter shared blocks from pairs that share less recent ancestry. Both estimates are consistent with a founder event in the range of ∼4,500–10,000 years ago, placing the origin of this *BRCA1* pathogenic variant in the Neolithic to Bronze Age period. The wide confidence interval (95% CI: 6,764–17,357 years) reflects the limited statistical power of n=7 carriers. With additional samples, the estimate would narrow considerably.

Taken together, the simple decay estimate (∼4,474 years) likely represents a lower bound, while the pairwise mean estimate (∼9,831 years) may be inflated by pairs with short, shared blocks due to phasing errors in the small sample. The true founder age is most likely in the range of 4,000–10,000 years ago.

## Discussion

These results support that c.4065_4068del (p.Asn1355Lysfs*10) represents a founder mutation in Rwanda. Across multiple datasets and populations, *BRCA1* c.4065_4068del (p.Asn1355Lysfs*10) emerges as a clearly pathogenic, recurrent frameshift variant, but current evidence does not yet establish a single, well-characterized founder origin. The deletion removes four nucleotides in exon 10, producing a frameshift at codon 1355 and a premature stop 10 amino acids downstream, and is consistently classified as pathogenic in ClinVar and clinical knowledgebases. It is very rare in general-population datasets (only a handful of alleles in gnomAD), yet it has been observed repeatedly in high-risk breast and ovarian cancer families and unselected cases from Europe, Asia, and the Middle East. In early European series, c.4065_4068del (often referred to as 4184del4) was among the common *BRCA1* mutations in high-risk breast/ovarian families from northwest England^15^, where haplotype analysis suggested that this and another recurrent mutation likely arose on specific ancestral chromosomes shared by multiple families, consistent with founder effects at regional or national scales. Large German and North American cohorts similarly reported this variant among recurrent *BRCA1* mutations, but without definitive evidence for a single pan-European founder^4, 16^.

More recent work extends this pattern to Asian and Middle Eastern populations^7–8, 18^ and again supports recurrence, with hints—but not definitive proof—of region-specific founder events. In a large Chinese ovarian cancer series (n=530), c.5470_5477del, c.981_982del, and c.4065_4068del were the three most frequent BRCA1 pathogenic variants, strongly suggesting that c.4065_4068del has been established in East Asian populations as a recurrent, and possibly founder, allele^17^. A Pakistani database of *BRCA1/2* variants found c.4065_4068del among 23 recurrent *BRCA1* mutations, and chased its origin alongside other frequent alleles, again pointing to a shared haplotype within that setting^8^. In the Gulf region, both a UAE series and an Emirati hereditary breast/ovarian cancer cohort identified c.4065_4068del as one of the most common *BRCA1* pathogenic variants, including four unrelated Emirati carriers in one recent clinic-based study^7, 18–19^. Laitman et al.’s regional synthesis of *BRCA1/2* PVs across Middle Eastern, North African, and South European countries shows that c.4065_4068del appears repeatedly in multiple countries, though it does not reach the very high, multi-country frequencies of “classical” founder variants such as c.68_69del, c.181T>G, or c.5266dup^9^.

Taken together, these reports indicate that *BRCA1* c.4065_4068del is a globally recurrent, clearly deleterious allele that has reached appreciable frequencies in several geographically and ancestrally distinct populations—including northwest Europe^15, 22^, Germany^4^, Pakistan^8^, China^17^, and the Arabian Gulf^7, 18^ —rather than being confined to a single, well-defined founder population. Several lines of evidence suggest that at least some of these clusters are due to local founder effects: (i) enrichment among high-risk families within specific regions (e.g., northwest England, Pakistan, parts of China), (ii) repeated identification in unrelated Emirati and UAE families, and (iii) the presence of shared extended haplotypes around *BRCA1* 4184del4 in earlier European work^23–24^. However, formal haplotype and coalescent analyses for c.4065_4068del have been limited and region-specific; no study has yet combined data across continents to infer a single ancestral origin or reconstruct migration paths. The available evidence is compatible with either a relatively old mutation that has spread and drifted in frequency across Eurasia and North Africa, or multiple, independent mutational events at a short deletion-prone motif.

Our Rwandan observations add a new and important dimension to this literature. Finding c.4065_4068del in 7 of 175 (4%) unselected breast cancer cases—and in 22% of all carriers of pathogenic variants—among women without reported close relatedness is striking compared with its rarity in global reference datasets and its generally modest frequencies in other series. Given historical links between East/Central Africa, the Middle East, and the Indian Ocean trading network^18–20^, one plausible hypothesis is that the Rwandan allele shares ancestry with a Middle Eastern or South Asian founder lineage (e.g., Pakistani or Emirati), but this cannot be assumed from frequency data alone. At present, no published study has performed cross-regional haplotype comparison of African, Middle Eastern, and Asian carriers of c.4065_4068del. Thus, while prior work strongly supports the notion that c.4065_4068del can behave as a founder variant within specific populations (northwest England, parts of Pakistan, China, and the Gulf), the geographic origin of the variant remains unresolved, and its presence in Rwanda could reflect either an independent founder event or introduction via historical gene flow from Middle Eastern or North African sources.

These gaps underscore the need for targeted haplotype and ancestry analyses of c.4065_4068del carriers across diverse populations. In our Rwandan series, high local frequency among apparently unrelated cases strongly suggests a founder effect in this population. By comparing extended haplotypes around *BRCA1* in Rwandan carriers with those reported (or that can be imputed) from European, Middle Eastern, North African, South Asian, and East Asian carriers, future work can test whether the Rwandan founder shares a common ancestral chromosome with any of these groups or represents a distinct African origin. Such analyses would not only clarify the evolutionary history of this clinically important variant but also inform region-specific genetic testing strategies and cascade screening in Rwanda and neighboring countries.

Our haplotype analyses suggest that the *BRCA1* c.4065_4068del (p.Asn1355Lysfs*10) pathogenic variant is an ancient founder allele in the Rwandan population, with an origin most plausibly between roughly 4,000 and 10,000 years ago. Using a simple decay-based approach applied to the longest haplotype segment shared by all seven carriers (581 kb; 0.3129 cM), we estimated a lower-bound age of approximately 160 generations, or ∼4,474 years (∼2,450 BCE, assuming 28 years per generation). This estimate reflects the minimum time required for recombination to erode a shared ancestral chromosome segment of this length and is constrained by the requirement that the segment be common to all carriers, which biases toward younger, more recent shared ancestry. A complementary pairwise approach that considered all 21 carrier pairs, allowing for a small mismatch tolerance to accommodate phasing error and local haplotype noise, yielded shorter average shared segments (mean 0.1424 cM; median 0.0319 cM) and a correspondingly older inferred age. Under this model, the pathogenic variant was estimated to have arisen around 351 generations ago (mean ∼9,831 years), with a wide 95% confidence interval of 242–620 generations (∼6,764–17,357 years). This older estimate reflects the inclusion of pairs who share only short residual haplotypes around *BRCA1*, which may arise from deeper ancestry or from methodological noise in a small sample. Importantly, both methods are directionally concordant in placing the origin of the variant well before recent historical times, in the late Neolithic to Bronze Age period, rather than arising from a recent demographic event such as the last few centuries of migration or admixture.

From a population-genetic and clinical perspective, these age estimates have several implications. First, they support a true founder effect rather than a de novo or very recent introduction: the variant has persisted through many generations of demographic change, recombination, and potential population bottlenecks, yet remains detectable at appreciable frequency (4% of tested breast cancer cases) in contemporary Rwanda. Second, an origin several thousand years ago is consistent with the possibility that c.4065_4068del may also be present—at low but non-zero frequency—in neighboring populations that share deep ancestry with Rwandans, even if it has not yet been widely reported in those settings. Third, the lack of a tightly constrained point estimate and the broad confidence interval highlight the limitations of mutation-age inference from only seven carriers; additional Rwandan and regional carriers, combined with higher-density phasing and cross-population haplotype comparison, will be needed to refine both the age estimate and any inference about whether the Rwandan founder shares a common origin with reported carriers in Europe, Asia, or the Middle East.

Overall, the most robust interpretation is that *BRCA1* c.4065_4068del is an old, likely pre-historical founder variant in this population, with a true age probably between ∼4,000 and 10,000 years. This time frame implies that the allele has been segregating through many generations of ancestry and social structure in the Great Lakes region. For clinical genetics and public health, this strengthens the rationale for including c.4065_4068del in targeted or low-cost testing panels for Rwanda and potentially for neighboring countries, and it motivates further work to map its geographical distribution, refine its ancestral origin, and characterize its penetrance and phenotype spectrum in diverse African settings.

Our previous findings suggest that this variant confers disease aggressiveness based on staging and disease features at presentation, compounded with the triple negative nature of the disease. Elsewhere, this variant has been associated with triple negative disease subtype^11, 18^, although stage and disease grade were not specified. In our parent study, the variant was not found among patients with prostate cancer.

In conclusion, our analysis reveals that the *BRCA1* c.4065_4068del variant is a pre-historical founder allele that has segregated within the Rwandan population from the late Neolithic or Bronze age, rather than being a result of recent gene flow. Further cross-regional studies are needed to determine if this lineage shares a common ancestor with global clusters or represents a distinct African origin, potentially refining our understanding of hereditary cancer risks across the continent.

## Declarations

### Ethical Approval and Consent to Participate

The seven (7) individuals were consented for additional testing and publication after the primary analysis^1^, as allowed by the larger Institutional Review Board approval (996/RNEC/2021 and 456/RNEC/2022).

## Consent for publication

The study participants consented for the data to be published

## Availability of supporting data

The whole-exome sequencing data generated in this study are not publicly available due to patient privacy and consent restrictions, but are available from the corresponding author upon reasonable request and subject to IRB approval. Clinical summary data are presented in the manuscript tables. Reference datasets from the 1000 Genomes Project (Phase 3, GRCh38) are publicly available at https://www.internationalgenome.org. The analytical code used in this study is available upon request.

## Authors’ contributions

AVCM, SB, TRR wrote the main manuscript AVCM, SB, TRR prepared the figures AMON, DK prepared figure 1

All authors reviewed and approved the manuscript

## Acknowledgements

N/A

